# Probing recombinant AAV capsid integrity and genome release under thermal stress by single-molecule interferometric scattering microscopy

**DOI:** 10.1101/2024.03.07.583968

**Authors:** Eduard H.T.M. Ebberink, Alisa Ruisinger, Markus Nuebel, Helena Meyer-Berg, Irene R.S. Ferreira, Marco Thomann, Albert J.R. Heck

## Abstract

Adeno-associated viruses (AAVs) are gaining traction as delivery vehicles for gene therapy although the molecular understanding of AAV-transgene release is still limited. Typically, the process of viral uncoating is investigated (*in vitro*) through thermal stress, revealing capsid disintegration at elevated temperatures. Here, we used single-molecule interferometric scattering microscopy to assess the (in)stability of different empty and filled AAV preparations. By introducing a heat-stable DNA plasmid as an internal standard, we quantitatively probed the impact of heat on AAVs. Generally, empty AAVs exhibited greater heat resistance than genome-filled particles. Our data also indicate that upon DNA release, the capsids do not transform into empty AAVs, but seem to aggregate or disintegrate. Strikingly, some AAVs exhibited an intermediate state with disrupted capsids but preserved bound genome, a feature that experimentally only emerged following incubation with a nuclease. Our data demonstrate that the thermal uncoating process is highly AAV specific (*i*.*e*., can be influenced by serotype, genome, host system). We argue that nuclease treatment in combination with mass photometry can be used as an additional analytical tool for assessing structural integrity of recombinant and/or clinical AAV vectors.

## Introduction

As a member of the *Dependoparvovirus* genus, adeno-associated viruses (AAVs) are not infectious by themselves and lack replication capabilities. Nevertheless, AAVs can enter mammalian cells, deliver their genetic cargo to the nucleus and transduce the cell, making them ideal candidates for gene therapy. Due to their non-pathogenic nature, low immunogenicity, and ability to sustain long-term expression recombinant AAV vectors (for the remaining document also referred to as “AAV”) are widely explored, with currently already half a dozen approved AAV-based gene therapies presented.^1-3^ However, this therapeutic approach is still relatively new and exhibits still several challenges. For instance, achieving high yield production with precise control over AAV packaging remains difficult, and the potential for unintended packaging of by-products persists. Additionally, the precise fate of AAV capsids upon host cell entry and factors affecting stability and transduction efficiency remain largely unclear.

Like other members of the *Parvoviridae* family, AAVs form small (∼25 nm diameter), icosahedral protein shells consisting of 60 capsid proteins (VPs). The capsid is built up from three different capsid protein isoforms (VP1, VP2 and VP3), differing mainly in the size of their N-terminal sequence with VP1 being larger than VP2 and VP2 being larger than VP3. The VP1:VP2:VP3 ratio is regarded to be in a 5:5:50 to 10:10:40 range although the exact stoichiometry is highly variable.^4,5^ Inside the capsid, AAVs can encapsulate a single-stranded DNA (ssDNA) transgene ideally limited to the size of the wildtype genome of about 4.8 kb. Due to the low transduction efficiency and broad tropism displayed by AAVs, current research is highly focused on improving AAV targeting and potency by rational design of both the capsid and transgene.

Despite ongoing research to improve AAV efficacy and furthermore establishing them as functional gene delivery tools for therapeutic applications, the understanding of capsid trafficking and transgene release remains incomplete. According to the current model of AAV transduction, capsids escape into the cytosol after endosomal uptake and become transported into the nucleus as intact capsids.^6-8^ It is within the nucleus that the ssDNA becomes accessible for further processing.^9^ An important step in the uncoating of AAVs is that the extended N-termini of VP1 and VP2 that normally reside within the capsid, emerge outward just before endosomal escape, in a likely pH-triggered event.^10-13^ This initial step then uncovers different nuclear localization sequences which are vital for nuclear uptake of AAVs either by the nuclear pore complex or *via* pore formation in the nuclear envelope.^6,8^ Once inside the nucleus, however, the process of AAV uncoating and ssDNA release remains elusive.

As investigating the behavior of AAVs within nuclei poses major challenges, *in vitro* experiments have been applied to simulate the AAV uncoating process, mostly induced by thermal energy. One important characteristic of AAV capsids is their high heat stability that reportedly allows them to endure temperatures up to 85 °C, at least for some serotypes.^14,15^ However, similar to cellular uptake, prolonged heating exposes the N-terminus of VP1/VP2 towards the capsid surface, and, eventually leads to complete uncoating of the ssDNA.^16-19^ To assess genome accessibility following heating of AAVs and similar parvoviruses (e.g. MVM, B19), techniques like electron microscopy (EM), atomic force microscopy, and analytical ultracentrifugation have been employed in combination with assessments of downstream capsid functionality, such as transduction efficiency, titer determination, and response to DNase treatment.^20-23^ Recent studies utilizing biophysical techniques like charge-detection mass spectrometry (CDMS) and mass photometry have also demonstrated that transgene size, pH and ionic strength can influence the thermal stability of AAVs.^24-28^ Yet, the intricacy and the significance between these different factors lack clarity and sometimes even appear contradictory (for instance in the effect of genome size).^15,19,24,25^ Thus far, two primary paths for AAV capsid uncoating have been proposed: one involving ssDNA externalization without capsid disassembly, or ssDNA externalization with complete dismantling of the capsid exterior.^21,23^ Nevertheless, a unified and comprehensive model elucidating the precise order of AAV uncoating either by thermal energy or within a cellular nucleus is lacking.

Characterization and quantification of AAV thermal uncoating by most of the above-mentioned techniques can be difficult as measurements are done under harsh circumstances (*e*.*g*. in the gas phase or by flash freezing), require laborious data analysis and/or lack an internal standard. Therefore, we adhere to single-molecule interferometric scattering microscopy measurements (also known as mass photometry (MP))^29-32^ with the capsids in solution under buffered conditions (*e*.*g*. PBS) and fast data acquisition to probe the effect of thermal energy on AAVs. This allows us to explore different assay set-ups and furthermore clarify and quantify ssDNA uncoating in reasonable throughput and sensitivity, allowing us to monitor the heating process in different (empty and filled) AAV serotypes and batches.

## Results

### Monitoring thermal AAV uncoating by mass photometry (MP)

AAV8 capsids produced by a HEK293-derived cell line and packaged with a CMV-GFP transgene (∼1 MDa in size) were measured by MP. These AAV8 capsids displayed a similar capsid distribution as previously reported for AAV8s from the same production platform (i.e., Revvity Gene Delivery formerly Sirion Biotech, termed AAV8_Rev_GFP throughout this paper).^33^ More than 90% of the capsid particles contain a transgene, and their Mw distribution is centered around ∼5 MDa (4.9 ± 0.08 MDa). Only a small population of about 8.4% is detected as empty particles (Mw ∼3.9 ± 0.05 MDa) (Fig. 1). When sampling the same AAVs following heat-treatment at 65 °C, a relative decrease in filled particles is observed with an apparent increase in the number of empty capsids (Fig. 1A). This heat-induced behavior is seemingly in line with previous CDMS and MP studies.^24-27^ To quantify the relative peak abundance three independent heating experiments were performed, normalized to the most abundant AAV peak (Fig. 1B). Following 5 minutes of heating, the population of filled particles decreases by a third to 61.4 ± 2.8 %. Prolonged heating furthermore drops the filled population to 42.2 ± 5.7 % with an accompanied increase in the empty part to 57.8 ± 5.7 % (Fig. 1C). While exposed to heat, a new peak in the MP mass histograms emerges centered at ∼1MDa (Fig. 1A). This mass corresponds nicely to that of an intact single ssDNA genome. Simultaneously, a rise in low molecular weight particles can be seen indicative of capsid disassembly. These initial data showed that incubation of AAV8_Rev_GFP at 65 °C steadily uncoats the ssDNA, leading to a loss of filled particles. In terms of absolute particle counts, however, only a slight increase of empty capsids can be observed when compared to the decrease in filled particles. This raises the question whether new empty capsids are formed as a consequence of the filled particles losing their genome (Fig. 1D), following a model proposed in recent studies.^24-27^ We argued that such an analysis would require an internal standard for qualitative and quantitative assessment of the number of particles in each sub-populations, ideally a high molecular weight standard that is insensitive to heat-treatment.

**Figure 1:**
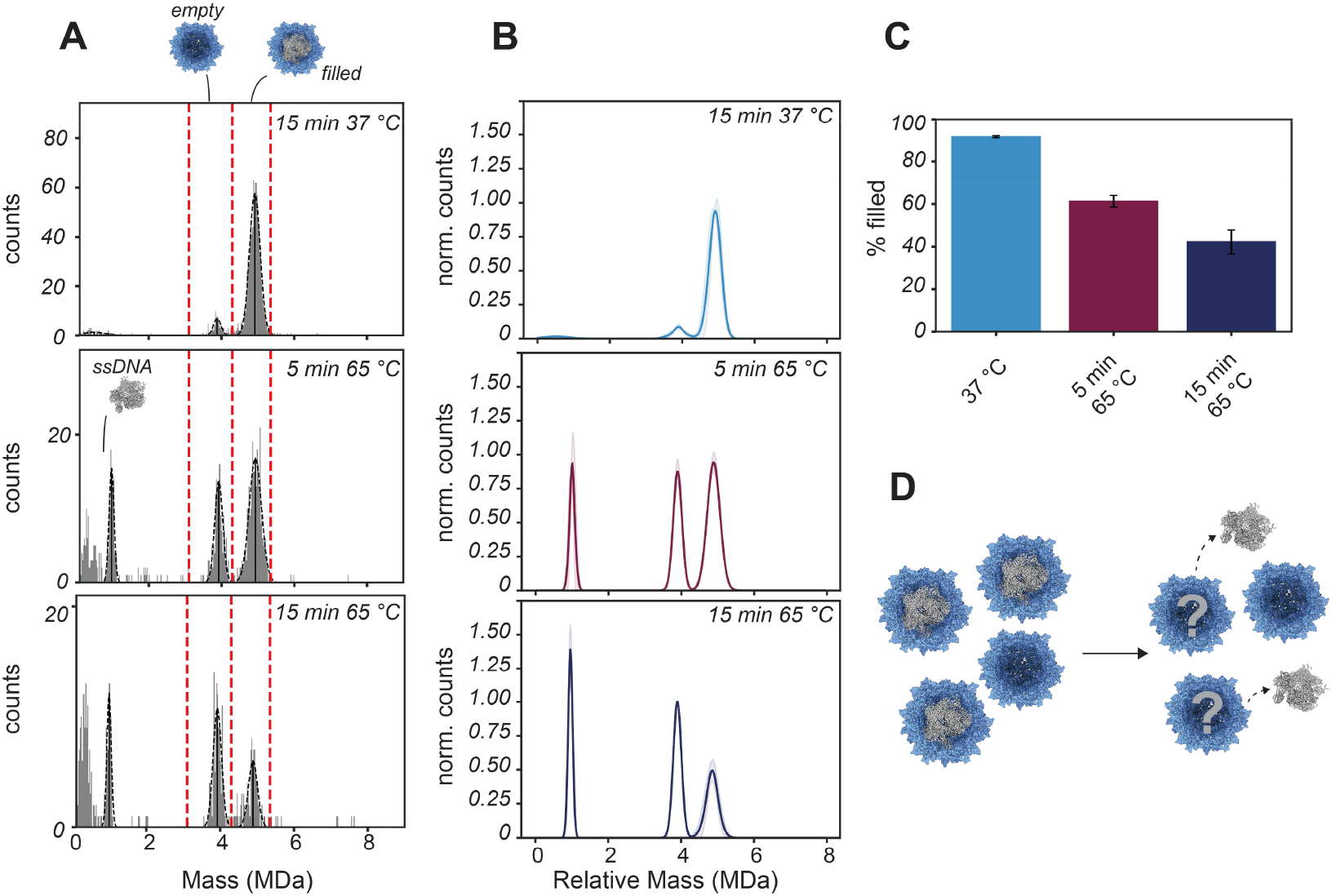
Uncoating of AAV8 capsids produced by a HEK293-derived cell line (AAV8_Rev_GFP) monitored by mass photometry. **A)** Raw mass histograms derived from the MP recordings. Before heat treatment, AAV8_Rev_GFP consists of approximately 92% filled particles and 8% empty particles. Capsids were incubated at either 37 °C or 65 °C, for either 5 minutes or 15 minutes, prior to analysis by MP. Prolonged incubation at 65 °C strongly decreased the filled population and gave rise to a new set of particles close to 1 MDa, a mass consistent with the mass of the released genome. **B)** Average Gaussian fits obtained from mass histograms of at least 3 MP repeats normalized by their counts. Counts were normalized to either the empty or filled particle population depending on which was most abundant, and the fits were aligned by their empty population. The standard deviation is given as semi-transparent bands. **C)** The percentage of filled capsids (% filled) decreases upon heat treatment. The mass range for quantification of particles is given in red dashed vertical lines in panel A (*i*.*e*., for empty particles between 3.1 and 4.3 MDa, for filled between 4.3 and 5.35 MDa). Average percentages are given with standard deviation extracted from measurements obtained in panel B. **D)** By monitoring the AAVs following heating at 65 °C, the release of the ssDNA becomes apparent, whereas it seems that relative to empty particles the filled AAVs disappear more strongly. Whether empty capsids are formed by release of the genome remains an open question.

### Use of the pBR322 plasmid as heat-stable, internal standard for mass photometry

To quantify the number of particles more accurately in each of the sub-populations co-appearing in the thermal uncoating of AAV capsids we next applied an internal quantitative standard, inert to heat treatment, at least up to 65 °C. Of note, we use this standard not for mass calibration, but solely for particle quantification. Because of the stable properties of dsDNA plasmids we chose the well-characterized plasmid pBR322 containing 4361 bp.^34,35^ As previously demonstrated, the for MP normally used standard, unmodified glass surfaces are not ideal for binding of dsDNA, so the coverslips were coated with (3-Aminopropyl)triethoxysilane (APTES) to detect the pBR322 landing events (Fig. 2A).^36^ On the APTES coated surface, dsDNA plasmids can be detected and analyzed by MP giving numerous contrasts values. However, to convert pBR322 contrast values into mass, a dsDNA based calibration is required.^36^ In our case, using a protein based calibrant, the MP measured pBR322 mass appears in the range of ∼1.8 MDa. The apparent mass of the pBR322 plasmid signal will not interfere with empty or filled AAVs and not with possible VP fragments of uncoated AAVs and thus has an ideal contrast value as reference for monitoring empty and filled AAVs. Notably, for pBR322 particles a subset of landing events displays oval-shaped signals instead of the anticipated circular shape based on the point-spread function (Fig. 2A). Such shapes most likely stem from supercoiled pBR322 plasmids that in length are larger than the diffraction limit tractable by the mass photometer.^36^ Heating the pBR322 sample at 65 °C for 15 minutes does not seem to influence the distribution of pBR322 landing events either in ellipticity or on the amount of analyzed particles and indicates a neglectable change of 1.04 ± 0.1 following heating (Fig. 2B). This confirmed that pBR322 can be used as quantitative reference standard in single-particle MP measurements during heating experiments.

**Figure 2:**
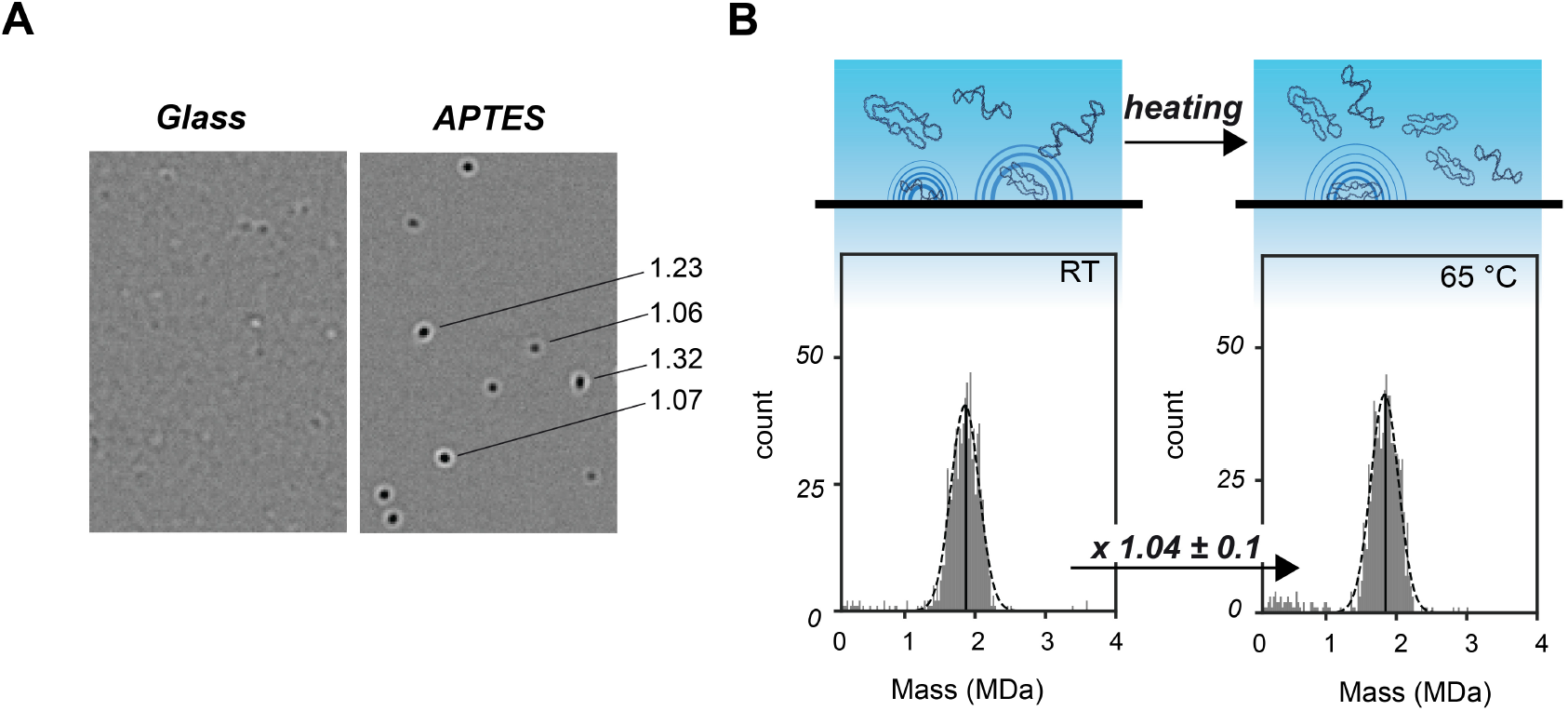
The use of pBR322 plasmid as quantitative reference standard in mass photometry. **A)** To capture landing events of pBR322 slides were coated with APTES enhancing the binding interaction. When detecting pBR322 some signals appear as ellipsis as also indicated by their elevated ellipticity value (=ellipse_width_ /ellipse_height_) seen on the right. **B)** After exclusion of oval-shaped landing events that do not fit the PSF, contrast value of pBR322 landing events were processed based on a thyroglobulin protein standard to produce a mass value. Resulting masses were binned in a mass histogram and fitted with a Gaussian curve (dashed, black lines) giving an average mass of about 1.8 MDa. In this representative measurement, heating at 65 °C for 15 minutes did not affect the number of particles measured (798 at RT versus 779 at 65 °C). Over three repeats, the change in landing events due to heating remained within 5%.

### Quantification of heated AAVs by mass photometry using the pBR322 plasmid as heat-stable internal standard

With the pBR332 reference plasmid spiked in the AAV8_Rev_GFP sample, we incubated these mixtures again at 65 °C for 15 minutes (Fig. 3A). As expected, this incubation did not deteriorate the pBR322 MP count. In contrast, the number of filled AAVs particles were found to be strongly reduced (Fig. 3A). Using the pBR322 signal we can normalize the AAV signals prior to heating and after heating. This way, we establish a near 40-fold decrease of filled AAV particles following heating at 65 °C (Fig. 3A). As seen before, the AAV8_Rev_GFP contains a small subset of empty AAV particles (∼8.5 %). Also, this particle population appears to slightly decrease, albeit by just 3-fold. The same approach of heating the AAV sample with pBR322 spiked in the solution was repeated at different incubation temperatures (Fig. 3B & S2). Following normalization, the decline in filled AAV particles can readily be observed starting at around 55 °C with a major drop at 60 °C (∼7-fold on average) and even further decrease at 65 °C. In comparison, the smaller sub-population of empty AAVs, already present before heat treatment, does not substantially decrease or increase (Fig. 3B). Empty capsids have been reported to be more heat-stable.^26^ To verify that finding we repeated the measurement with empty AAV8 capsids, produced in HEK293-cells in the absence of a transfer plasmid. When analyzing this pool of AAV8_Rev_empty capsids, it confirmed that empty AAV8 capsids are relatively heat-stable, remaining largely unaffected by heat-stress up to 65 °C (Fig. 3C and S2). At higher temperatures T > 65 °C, however, these capsids also disintegrate (Fig. 3C). Heat treatment of the AAV8_Rev capsids shows that capsids filled with a ssDNA genome rapidly break down, while empty capsids remain more stable. Notably, and in contrast with what has been hypothesized in recent studies,^24,26,27^ the disappearance of filled capsids is not compensated by a concurrent increase in empty capsids.

**Figure 3:**
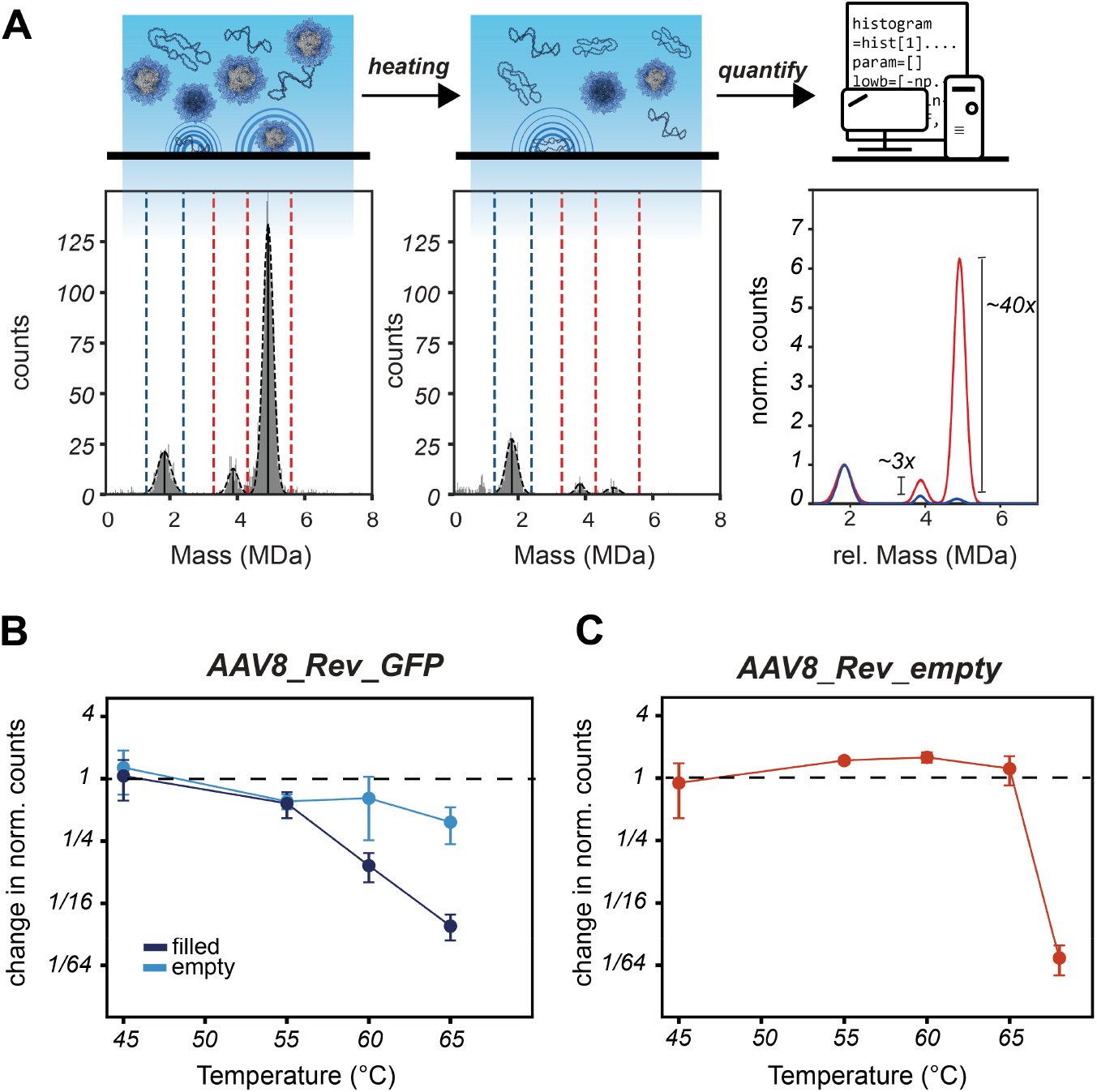
Monitoring the fate of AAV8 capsids produced by a HEK293-derived cell line upon heating using a heat-stable DNA plasmid as internal standard. **A)** Together with the pBR322 reference plasmid, AAV8_Rev_GFP capsids were incubated at 65 °C for 15 minutes and compared to the same sample prior to heating (RT, room temperature). The binned landing events measured on APTES slides were fitted with Gaussian curves displayed as dashed, black lines. To quantify the number of empty and filled capsids, using pBR322 as internal standard, different mass ranges were used as indicated by dashed vertical lines. Using pBR322, the AAV population was normalized, aligned and the change in abundance determined. **B)** Fold-change of the number of AAV capsids upon heating (for 15 min) compared to incubation at room temperature. When increasing the temperature, the filled capsids start to deteriorate, eventually at higher temperatures also followed by some loss of empty capsids. **C)** Shown is the stability of AAV8s that were prepared in absence of a genome (purposefully produced empty). Following heat treatment (15 min), the MP reveals that AAV8_Rev_empty is extremely stable till at least 65 °C. Error bars in panel B and C represent the standard deviation.

### The release of ssDNA genome proceeds via disordered intermediate states wherein the DNA is accessible and prone to hydrolysis

In the AAV8_Rev_GFP heating experiments, a novel population of particles emerged with a mass of approx. 1 MDa, likely originating from ssDNA released from the capsids (Fig. 1). To probe whether the suspected ssDNA population is sensitive to DNase degradation we heated AAV8_Rev_GFP at 65 °C for 15 minutes followed by incubation with DNase I and Mg^2+^ (Fig. 4A). After addition of DNase the peak at ∼1 MDa peak vanished, confirming that these particles are made up by DNA. Intriguingly, DNase treatment also resulted in a substantial drop in the number of AAV particles with masses of ∼5 MDa, initially assigned as intact, filled AAVs. The percentage of these apparent intact and genome containing AAVs before treatment with DNase was 54.3 ± 3.0 % (Fig. 4B). However, incubation with DNase decreased the percentage by a 1.7-fold to 31.2 ± 3.3 %. Triggered by this observation, we repeated the DNase treatments of AAV8_Rev_GFP, now incubated at different heating temperatures before adding the DNase (Fig. 4C). When keeping the AAV8_Rev_GFP capsids at room temperature the AAVs remain stable and insensitive to DNase. But when first incubated at temperatures above 45 °C, followed by DNase addition, part of the filled AAV population (assumption based on their mass) starts to disappear (Fig. 4C). At 55 °C and 65 °C the DNase driven decline in filled AAVs is noticeable, as reflected by a decrease in percentage (Fig. 4B and C). These findings demonstrate that we can detect a subset of AAV particles that are in an intermediate state of uncoating following heating, where the ssDNA genome is accessible, albeit still associated to the capsid.

**Figure 4:**
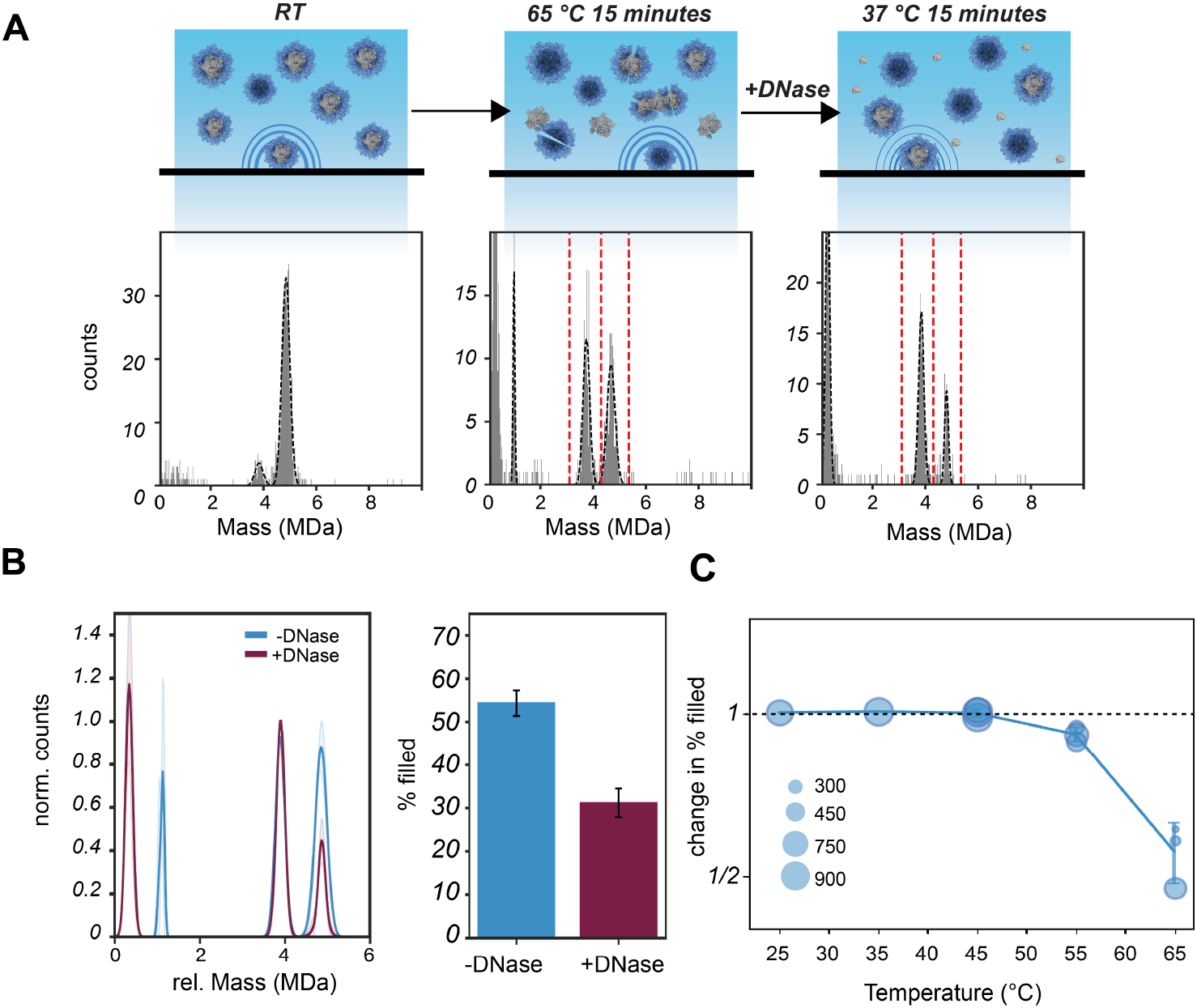
DNase induced change in the number of filled capsids of HEK293-derived AAV8. **A)** AAV8_Rev_GFP capsids were pre-heated at 65 °C, sampled by MP, treated with DNase I and again sampled by MP. Given are illustrative mass histograms which were fitted with Gaussian curves (black dashed curve). Following initial heating at 65 °C a new population of particles is detected with a mass of ∼1 MDa, presumably consisting of released ssDNA. It is evident that treatment with the DNase makes the released DNA peak vanish, but also leads to an elevated disappearance of filled particles, decreasing the measured percentage of filled capsids. Incubation with only Mg^2+^ did not display a decrease in filled particles (see Fig. S6). **B)** After repeating the procedure (n=3), an average Gaussian fit was constructed following normalization to the most abundant AAV peak within each repeat (standard deviation is given as semi-transparent bands). At the same time percentage of filled AAVs, represented here as bar-plots, were calculated based on the mass range indicated in the mass histograms by red dashed lines in panel A (AAV8_Rev_GFP empty: 3.1 - 4.3 MDa, AAV8_Rev_GFP filled: 4.3 - 5.35 MDa). **C)** DNase treatment was performed at several pre-heating temperatures. The acquired change in filled AAVs following DNase treatment is plotted as single circles: change in % filled=[% filled_+DNase_]/[% filled_-DNase_]. Because pre-heating induces a relative loss of filled AAVs, the amount of AAVs assessed before DNase treatment reduces with increasing temperature as depicted in the size of the circles. Error bars in panel B and C represent the standard deviation between the different repeats.

### Thermal uncoating of different AAV preparations

To assess whether the observed DNase induced degradation of pre-heated AAVs follows a general mechanism we next studied three AAV preps from different production platforms and serotypes as production method and serotype can strongly affect AAV properties.^37^ One of the AAVs we evaluated was an insect cell produced AAV8 containing a CMV_GFP transgene (0.7 MDa in genome size) acquired from Virovek, here termed sample AAV8_Vir_GFP (Fig. S1). As seen for the previous AAVs, incubation at 65 °C with subsequent addition of DNase gave a drop in apparent filled AAVs (Fig. 5A and S3). Notably, a small peak at approx. 0.7 MDa, at the mass of the genome, can be seen that also disappeared following DNase treatment (Fig. 5A). Here, the percentage of filled AAVs went from 42.0 ± 1.2 % to 20.3 ± 0.7%, a nearly 2-fold decrease, which gives the impression that the nuclease induced transgene degradation is even more pronounced for this AAV sample. Also, when monitoring the decline in filled AAVs at different pre-heating temperatures the onset of nuclease induced deterioration of the AAV8_Vir_GFP seems to occur earlier when compared to AAV8_Rev_GFP. Remarkably, even keeping the AAV8_Vir_GFP at 35 °C followed by nuclease addition gave a small but substantial drop in the filled AAV8_Vir_GFP particles.

**Figure 5:**
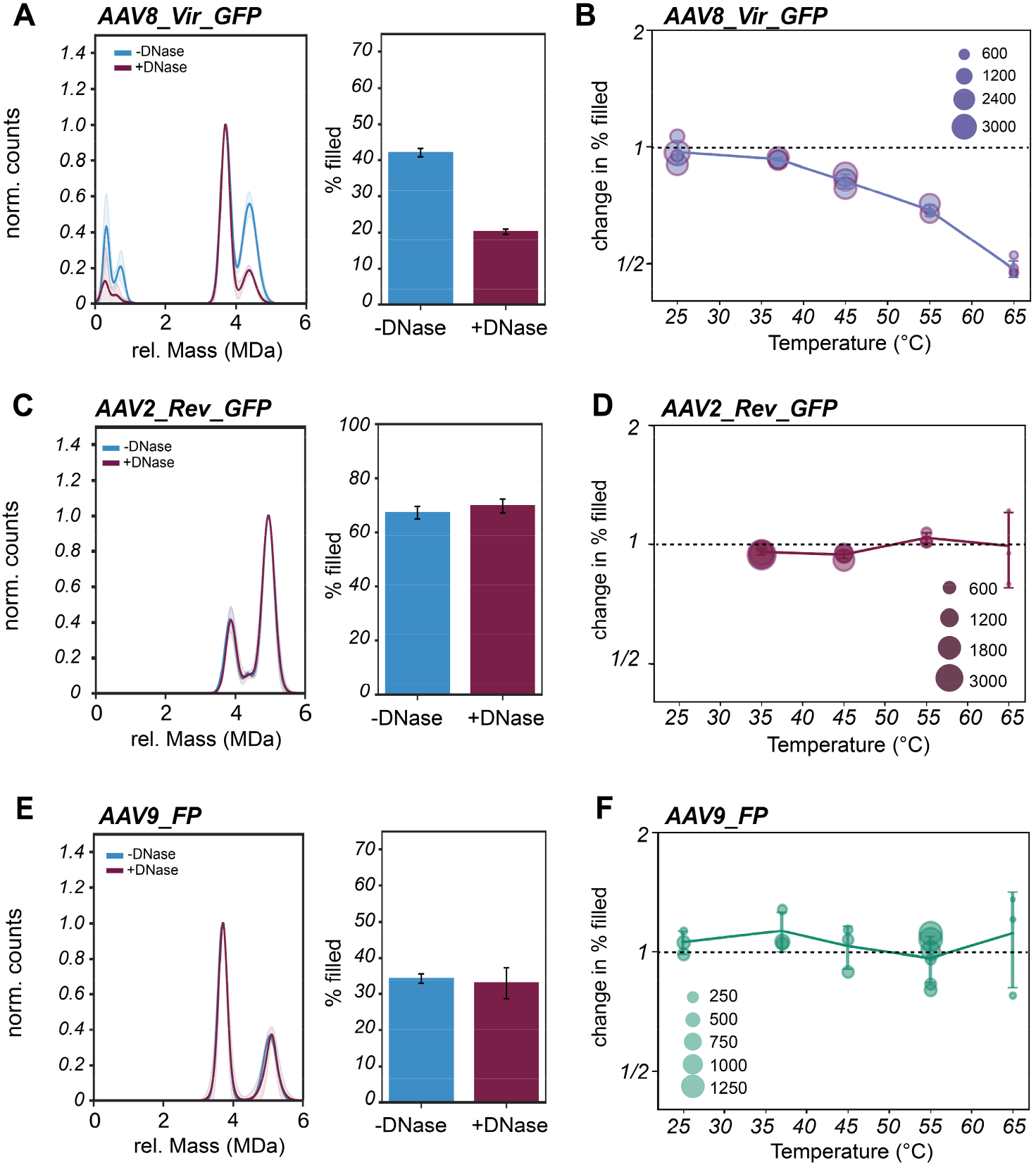
DNase induced changes in AAV8 from insect cells (AAV8_Vir_GFP), serotype AAV2 (AAV2_Rev_GFP) and serotype AAV9 (AAV9_FP) from HEK293 cells. AAV8s that were produced in insect cells, also containing an CMV-GFP genome were subjected to heating followed by DNase treatment. **A)** After incubation at 65 °C, DNase also induced a drop in the seemingly filled AAVs of AAV8_Vir_GFP. Given are the average Gaussian fits that followed from normalization to the most abundant AAV peak over at least three repeats. Quantification of the AAVs give a percentage of filled AAVs that decreased substantially after addition of DNase as can be seen in the bar-plot. **B)** The nuclease driven decline in filled AAVs begins already at relatively low temperatures compared to previously measured AAV8_Rev_GFP. For each measurement at different pre-heating temperatures, the change in % filled AAVs due to DNase incubation is given by a circle. The size of the circle indicates the amount of filled AAVs measured following pre-heating and before addition of DNase. The same experiment was done on AAV2 capsids containing a CMV-GFP transgene. **C)** Surprisingly, the AAV2_Rev_GFP capsids were not affected substantially to the nuclease treatment. Because at 65 °C nearly all AAV2 capsids are lost, we show DNase treatment of AAV2_Rev_GFP pre-heated at 55 °C. **D)** When plotting over different incubation temperatures no change in empty/filled AAV distribution can be observed for AAV2_Rev_GFP upon addition of DNase. **E)** Also, AAV9_FP capsids were analyzed using the same consecutive heating and nuclease incubation steps. Most capsids are lost at 65 °C, therefore, displayed are the average Gaussian fits of AAV9_FP that were heated at 55 °C and then incubated with DNase. **F)** When repeating over different incubation temperatures no substantial change in the percentage of filled AAVs can be observed for AAV9_FP. Error bars represent the standard deviation between the different repeats. The standard deviation in Gaussian fits is indicated by shaded, semi-transparent bands.

Next, we switched serotype and monitored AAV2 capsids that contain an identical transgene as AAV8_Rev_GFP (AAV2_Rev_GFP, supplied by Revvity Gene Delivery) and an AAV9 serotype with a fluorescent protein encoding transgene both produced from mammalian HEK293 cells (here termed AAV9_FP, with a 1.4 MDa genome size) (Fig. S1). When applying the 15-minute heating procedure followed by DNase treatment, the capsids of AAV2_Rev_GFP and AAV9_FP behaved distinctively from the AAV8 serotypes. After heating the capsids, AAV2_Rev_GFP did not seem to be affected at all by the subsequent nuclease treatment (Fig. 5C & D). Also, for AAV9_FP, incubation with DNase after heating did not change the number of filled particles, regardless of the pre-heating temperatures used (Fig. 5E & F). In addition to the lack of a DNase response, we could also not detect any obvious ssDNA landing events upon heating (Fig. 5C and E, Fig. S3 and Fig. S4). Of note, both AAV2_Rev_GFP and AAV9_FP, appear more prone to capsid disassembly as incubation at 65 °C leaves only a fraction of the initial number of filled AAV particles (Fig. S4). In addition, we observed substantial aggregation for the AAV2_Rev_GFP sample, which was not that apparent in the other studied samples (Fig. S5). Together, we observed distinctive behavior of the filled AAV9_FP and AAV2 capsids compared to the studied AAV8 capsids. Therefore, extracting a general mechanism of thermal uncoating of AAVs seems unfeasible.

## Discussion

AAVs have gained a pivotal role as vehicle in advanced gene therapies, and therefore AAVs are extensively studied *in vitro* and *in vivo*,^1,3^ with a focus on their production and function. However, the precise uncoating process of AAVs at the molecular level and subsequent release of the genetic cargo is still rather elusive. Stressing AAVs by thermal energy has extensively been used as a model to emulate the uncoating process and has recently seen new advancements by using techniques such as AFM, CDMS, and MP.^23-27^ Here, we employed MP in combination with a DNA plasmid-based reference standard to quantitatively assess the heat-induced uncoating process of AAVs. In our experiments AAV particles containing genomes disassembled under thermal stress, while the co-produced empty capsids remained largely unaffected (Fig. 3 & S4). Capsids that were produced without a transfer plasmid, and therefore unambiguous empty, even showed stronger stability. This confirms previous findings that empty AAVs exhibit greater heat-stability, while filled AAVs lose their integrity early upon heating.^26^ Therefore, generally, genome packing decreases the stability of the AAV capsids. In contrast to earlier propositions,^24,26,27^ we find here that when the number of filled AAVs decrease upon heating, we do not observe a substantial increase in empty AAV capsids (Fig. 3 & S4). This discrepancy with earlier results can perhaps be explained by the lack of a quantitative internal standard in these earlier experiments.

Interestingly, we were able to monitor by MP not only the fate of the AAV capsids but also, in several cases the formation of the released ssDNA (Fig. 1). For both AAV8_Rev_GFP and AAV8_Vir_GFP the released genomes could be detected, whereas for AAV2_Rev_GFP, which shares an identical transgene as AAV8_Rev_GFP, and for AAV9_FP detection of the release genome failed (Fig. 5). Likely, in these latter cases the released ssDNA co-aggregates with the disintegrating capsid. It is known that AAV capsids tend to aggregate upon heating,^27,38^ and our MP recordings revealed more aggregates upon heating. Especially, in the case of AAV2_Rev_GFP also at lower temperatures aggregates could already be detected (Fig. S5).

When we applied DNase to confirm the presence of the released ssDNA, we simultaneously observed a decrease in the number of apparently filled AAV8 capsids, as illustrated in Figure 4. This observation sparked our interest because intact filled AAVs are known to be resistant to nucleases.^21^ The loss of filled capsids initiated by the DNase treatment suggests the existence of an intermediate state where the transgene becomes exposed to the solvent while being retained to the partly disintegrated capsid. Such behavior has been described before by Bernaud et al., who used AFM to describe externalization of genomes without the disassembly of AAV capsids via a two-step ejection model.^23^ According to their measurements, the ssDNA genome remains connected to presumably intact capsids. A similar AAV state has also been documented by CDMS measurements.^24^ However, heating with the potential for genome ejection and release did not lead to the accumulation of stable, empty capsids (Fig. 3 and S4). Alternatively, heating may lead to a compromised capsid structure, triggering AAV disassembly. At this point, DNase can access and digest the genome, resulting in a loss of seemingly intact, filled AAVs (Fig. 4 & 5). In addition, the impaired capsids can potentially be captured in aggregates through ssDNA and VP connections as seen earlier.^19^ This results in a mixture of AAV capsid states either broken, aggregated, or emptied with or without transgene attached (Fig. 6). Our data reveal that depending on the serotype and incorporated transgene, potentially a (large) part of the capsids can be compromised this way. Incubation with DNase might equally disintegrate broken capsids as well as the aggregates.

**Figure 6.**
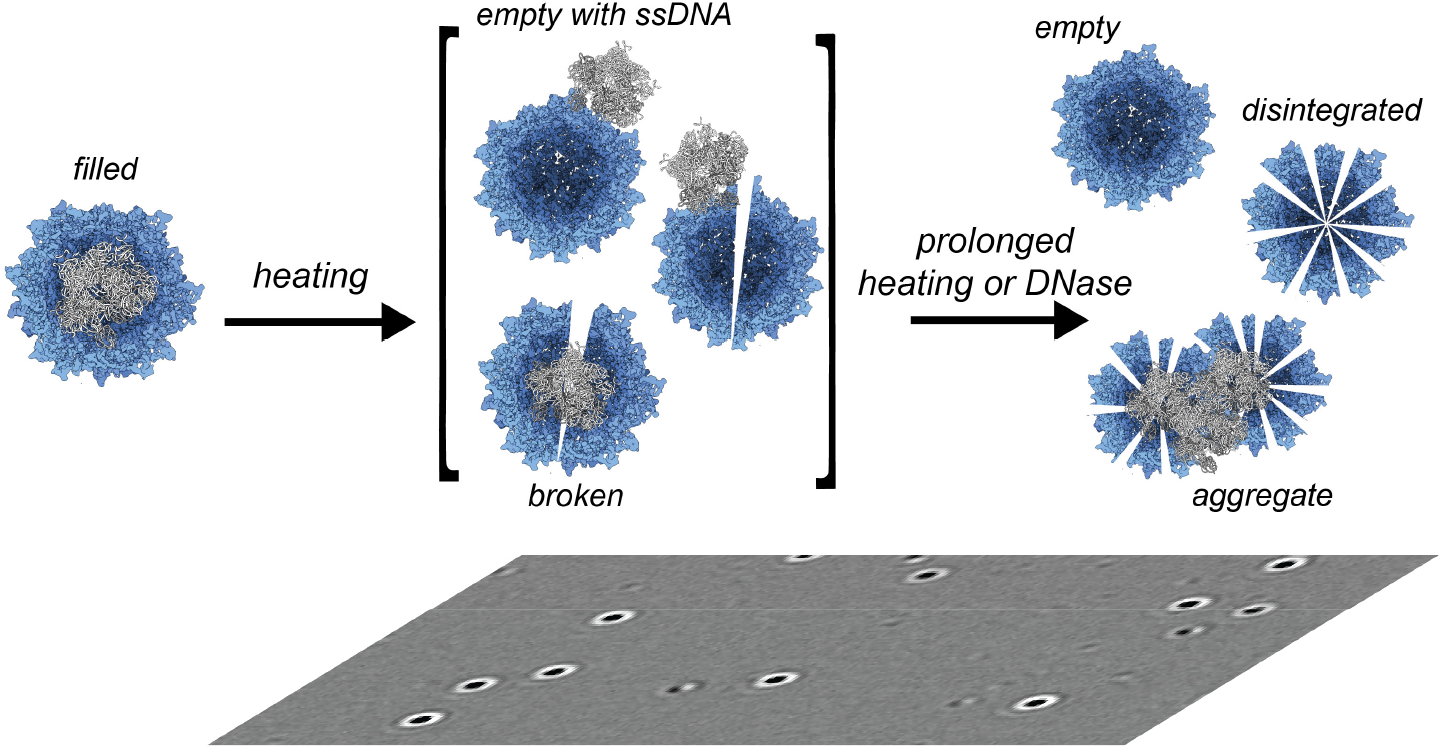
Model describing the fate of filled AAVs upon heating. Moderate heating (T < 60 °C) keeps empty AAV particles largely intact. However, genome filled particles start to disintegrate already between 55-65 °C. This process can lead to release of the ssDNA genome. Heating can also lead to distorted, partly open/broken AAV capsids to which the ssDNA is still attached having an indistinguishable mass when compared to the authentic encapsulated genome-filled AAVs. Further heating can lead to full disintegration of the capsid or, alternatively, to aggregation of broken capsids. DNase treatment can dissolute the attached ssDNA that is still attached to partly disintegrated AAV capsids. Still, the here presented data reveal also that each produced AAV may react differently to heating, with serotype, size of genome and production platform, being key factors.

The approach taken here, incorporating DNase treatment prior to MP measurements allowed us to uncover a previously elusive population of AAVs, that based on their mass could be wrongly assigned as intact filled AAVs. Notably, in the AAV8_Vir_GFP sample, this population was already present in AAVs incubated at relatively low temperatures (Fig. 5A). This implies that AAV damage can be induced easily, which here only became traceable by nuclease addition. To test AAV quality, an assay as presented here combining MP with DNase might be important, especially when considering that different AAV samples and/or production batches, and not just serotypes, display rather different behavior upon stress (*e*.*g*., heating) and DNase treatment. Moreover, the use of an internal standard, such as the pBR322 plasmid used here, in the quantitation of empty/filled/disintegrated AAV particles is highly recommended, when studying AAV stability.

## Materials and methods

### Preparation of (coated) coverslips for MP

Glass coverslips (Paul Marienfeld GmbH, 24 x 50 mm, 170 ± 5 μm) were overnight incubated in 100 mM Sulphuric acid (Merck). Afterwards the coverslips were rinsed consecutively with Milli-Q water, methanol (Biosolve Chimie SARL, HPLC grade), ethanol (Supelco EMSURE), methanol and finally left in ethanol. The ethanol was replaced with a mixture of 5% APTES (Sigma) in ethanol to coat the slides for 1 hour. The coated coverslips were rinsed twice with ethanol before incubation in 6% acetic acid (Merck) for 30 minutes. Afterwards slides were rinsed with methanol once, sonicated for 5 minutes in methanol and once more rinsed with methanol. Before usage, coverslips were rinsed with isopropanol (Supelco EMSURE) and dried with N_2_. Non-coated glass coverslips were prepared by serial rinsing with Milli-Q water and HPLC-grade isopropanol and subsequent drying with N_2_. Once dry, CultureWell gaskets (Grace Biolabs) were placed on the coverslips as container well for MP measurements.

### Mass photometry

All mass photometry measurements were executed by use of a Samux mass photometer (Refeyn Ltd.) and were performed following the same experimental procedure. Firstly, the mass photometer was allowed to focus by applying 12 μL of PBS buffer within a container well to either a glass or APTES coated coverslip mounted on the mass photometer. After focusing, each measurement was initiated by mixing of 3 μL of sample to the 12 μL of buffer prior to recording for 60 seconds with 100 frames per second. Contrast values were converted to mass values by measuring a thyroglobulin multimer mix (Sigma, T9145) and alignment of the three thyroglobulin contrast values to masses of 335, 670 and 1340 kDa. Mass values were processed by DiscoverMP software (Refeyn Ltd.) and exported for further processing by in-house prepared Python scripts. For MP characterization of pBR322 a sample was prepared by dilution of the pBR322 plasmid (Thermo Scientific, SD0041) in PBS buffer to a concentration of 125 ng/μL. The ellipticity of pBR322 landing events were calculated by extracting the outer contour of a landing event, fitting an ellipse, and dividing the width by its height.

### Mass photometry measurement of thermal stressed AAVs with pBR322 standard

The MP measurements with pBR322 were all carried out on APTES coated coverslips. Samples were prepared by mixing ∼125 ng/μL pBR322 and ∼1 x 10^12^ vg/mL of AAVs together in 10 μL PBS. A single MP measurement was done prior to heating (at room temperature) followed by heat incubation for 15 minutes at various temperatures in a thermomixer (Thermo Scientific). The heated sample was allowed to cool down to room temperature and was shortly spun down prior to the second MP measurement. To assess the effect of heating, counts were firstly quantified within specified mass ranges. For an equal assessment, mass histograms were aligned prior to quantification (*e*.*g*., to the pBR322 signal set at 1.8 MDa for AAV8_Rev_GFP). Secondly, the AAV counts were adjusted by the ratio between the pBR322 signal measured at room temperature and pBR322 signal measured at the elevated temperature (pBR322_RT_/pBR322_heated_). Finally, the relative change between room temperature and heated AAV populations was calculated by division of these pBR322 adjusted values.

### Mass photometry measurement of thermal stressed AAVs with DNase treatment

An amount of approx. 1 to 5 x 10^12^ vg/mL of AAVs in 15 μL PBS was heated for 15 minutes at various temperatures. Following heat incubation, a sample of the AAV solution was taken and measured by MP. Measurements were done on cleaned glass coverslips. Digestion of released and/or accessible DNA in the remaining AAV solution was initiated by addition of 100 nM of DNase I (Sigma, D5025) with subsequent addition of approx. 5 mM Mg Acetate (Sigma, M0631). The DNase/AAV mix was incubated for 15 minutes at 37 °C in a thermomixer. Following incubation with DNase, another sample was taken and measured by MP. For equal assessment amongst repeats the masses were aligned to the most abundant AAV population. Based on the quantified AAV counts, a percentage of filled was calculated (% filled=filled/[total AAV population]*100) of both the MP measurements done before and after addition of DNase (respectively % filled_-DNase_ and % filled_+DNase_). The relative change in % filled was given by division of both these percentages (*i*.*e*., change in % filled = [% filled_+DNase_]/[ % filled_-DNase_]).

## Data availability statement

No data availability.

## Acknowledgments

We thank the members of the Heck laboratory for general support. This research received funding by the Netherlands Organization for Scientific Research (NWO) through the Spinoza Award SPI.2017.028 to AJRH. This project received further support from Roche Diagnostics GmbH, Penzberg, Germany.

## Author contributions

EHTME conceptualized the project and performed all experiments and did the data analysis, wrote the first draft, and edited the final draft. AR, MN and MT supplied samples, provided funding and edited the manuscript. HMB and IRSF supplied (Revvity Gene Delivery) samples and edited the manuscript. AJRH conceptualized the project, provided supervision and financial support and infrastructure, co-wrote the first draft and edited the final draft.

## Declaration of interests

AR, MN and MT are employees of Roche Diagnostics GmbH, Penzberg, Germany, a company with interest in employing recombinant AAV vectors for gene delivery purposes. IRSF is an employee of Revvity Gene Delivery, Graefelfing, Germany, a company developing AAV vectors for gene delivery purposes. HMB was an employee of Revvity Gene Delivery and is an employee of Roche Diagnostics GmbH, Penzberg, Germany. The remaining authors declare no competing interests.

