## Supplementary Data for "Probing recombinant AAV capsid integrity and genome release under thermal stress by single-molecule interferometric scattering microscopy"

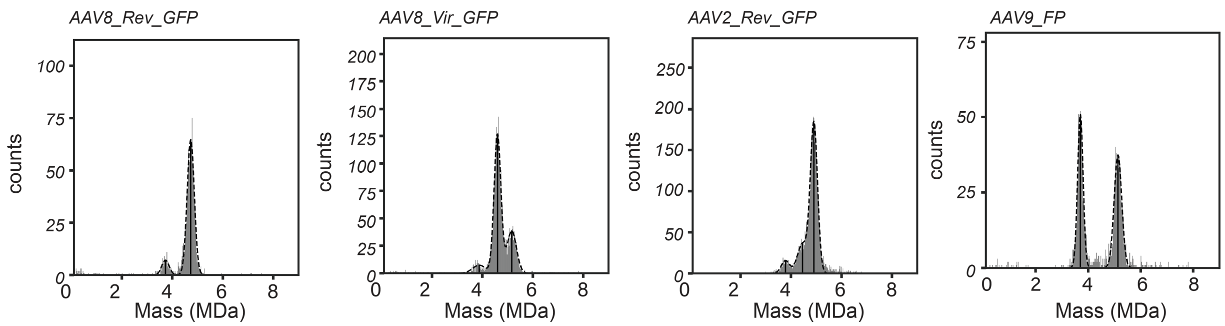


**Supplementary Figure 1:** ***Mass photometry measurement of the different AAV preparations AAV8_Rev_GFP, AAV8_Vir_GFP, AAV2_Rev_GFP and AAV9_FP.*** Mass histograms derived from MP measurements on the AAV preparations that were assessed in this study prior to any treatments with heat or a nuclease. The MP measurements shown here were done at room temperature in PBS and on glass slides. The AAV subpopulations were fitted with a Gaussian curve, indicated by black dashed lines. For every shown AAV preparation there is a considerable amount empty capsids present, especially in AAV9_FP. AAV8_Vir_GFP has capsids that contain more mass then expected based on a single genome (overfilled AAVs), while AAV2_Rev_GFP seems to have partially filled capsids.

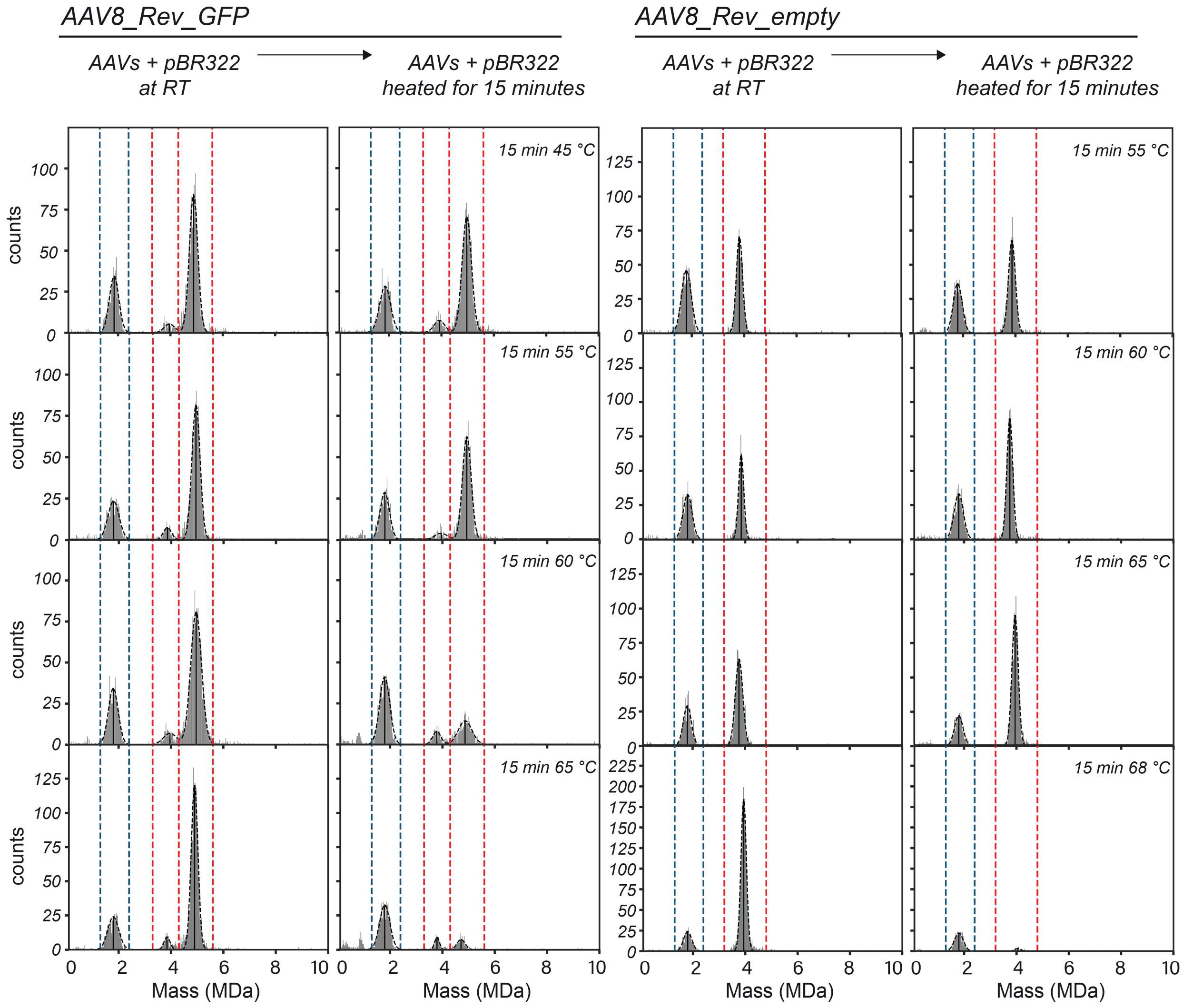


**Supplementary Figure 2:** ***Mass photometry quantification before and after heat incubation with pBR322 of AAV8_Rev_GFP and AAV8_Rev_empty.*** Illustrative mass histograms derived from MP measurements of the AAV8_Rev preparations (either with an CMV-GFP transgene or empty) that were heat incubated together with the pBR322 DNA plasmid for quantification. A loss of AAVs can be seen when comparing the amounts of AAVs measured at room temperature (RT, left side) against those that were heated (right side). Gaussian fits of the AAVs and pBR322 are indicated by black dashed lines. The mass ranges that were used for quantification of the different AAV subpopulations (see Figure 3) are indicated by red dashed lines, the mass ranges for pBR322 quantification are indicated by blue dashed lines (pBR322: 1.3 - 2.4 MDa, AAV8_Rev_GFP empty: 3.3 - 4.3 MDa, AAV8_Rev_GFP filled: 4.3 – 5.6 MDa, AAV8_Rev_empty: 3.2 – 4.8 MDa).

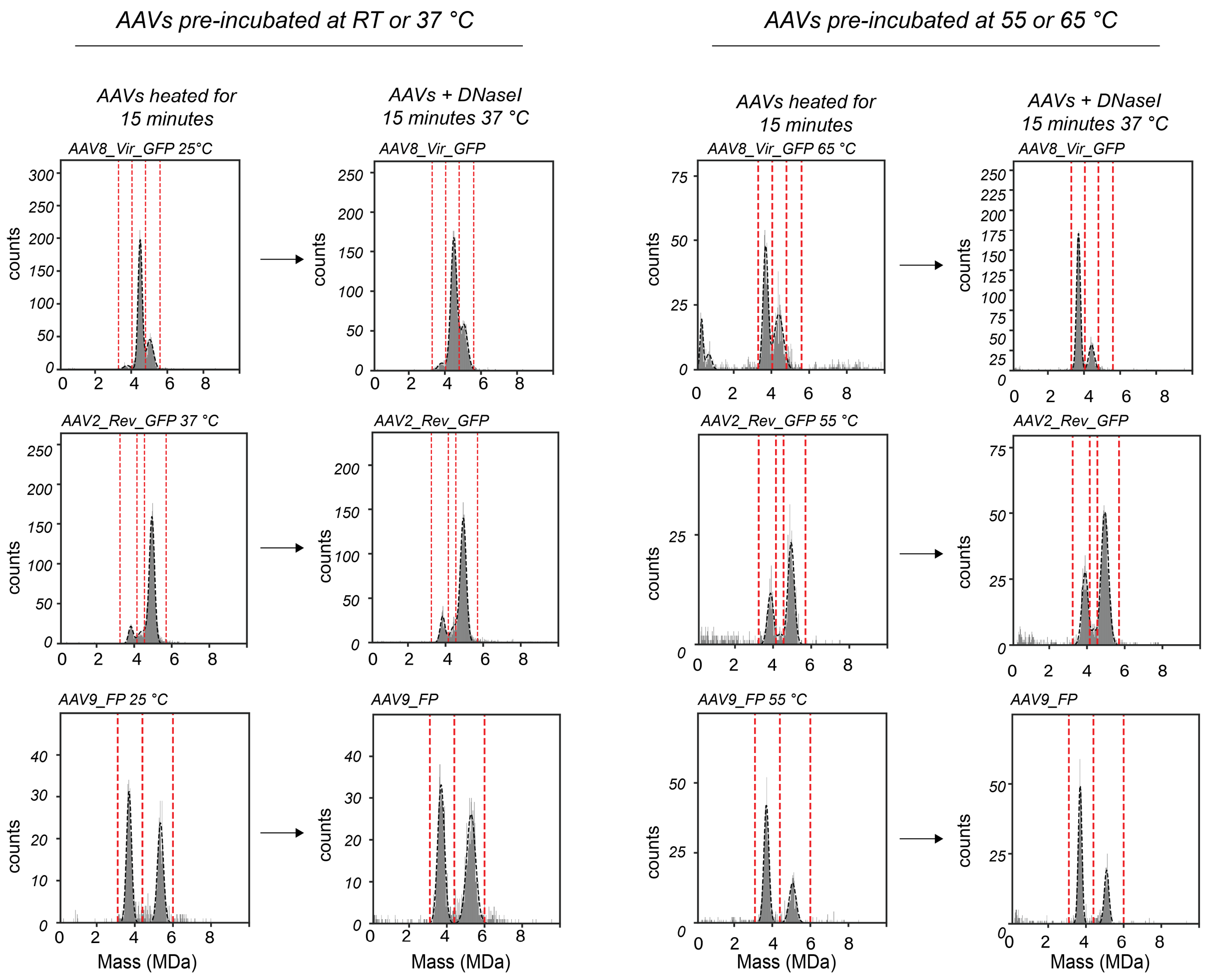


**Supplementary Figure 3:** ***Mass photometry quantification of pre-heated AAV8_Vir_GFP, AAV2_Rev_GFP and AAV9_FP before and after treatment with DNase.*** Displayed are representative mass histograms of AAV8_Vir_GFP, AAV2_Rev_GFP and AAV9_FP incubated at low or moderate temperatures (RT or 37 °C) or heated temperatures (55 or 65 °C) before and after addition of DNase. The different AAV distributions were fitted with a Gaussian curve indicated by black dashed lines. Mass ranges used for quantification of the AAV particles (see Figure 5) are indicated by red dashed lines (AAV8_Vir_GFP empty: 3.3 – 4.05 MDa, AAV8_Vir_GFP filled: 4.05 – 4.8 MDa, AAV8_Vir_GFP overfilled: 4.8 – 5.6 MDa, AAV2_Rev_GFP empty: 3.25 – 4.15 MDa, AAV2_Rev_GFP partially filled: 4.15 – 4.55 MDa, AAV2_Rev_GFP filled: 4.55 – 5.7 MDa, AAV9_FP empty: 3.1 – 4.4 MDa, AAV9_FP filled: 4.4 – 6.0 MDa).

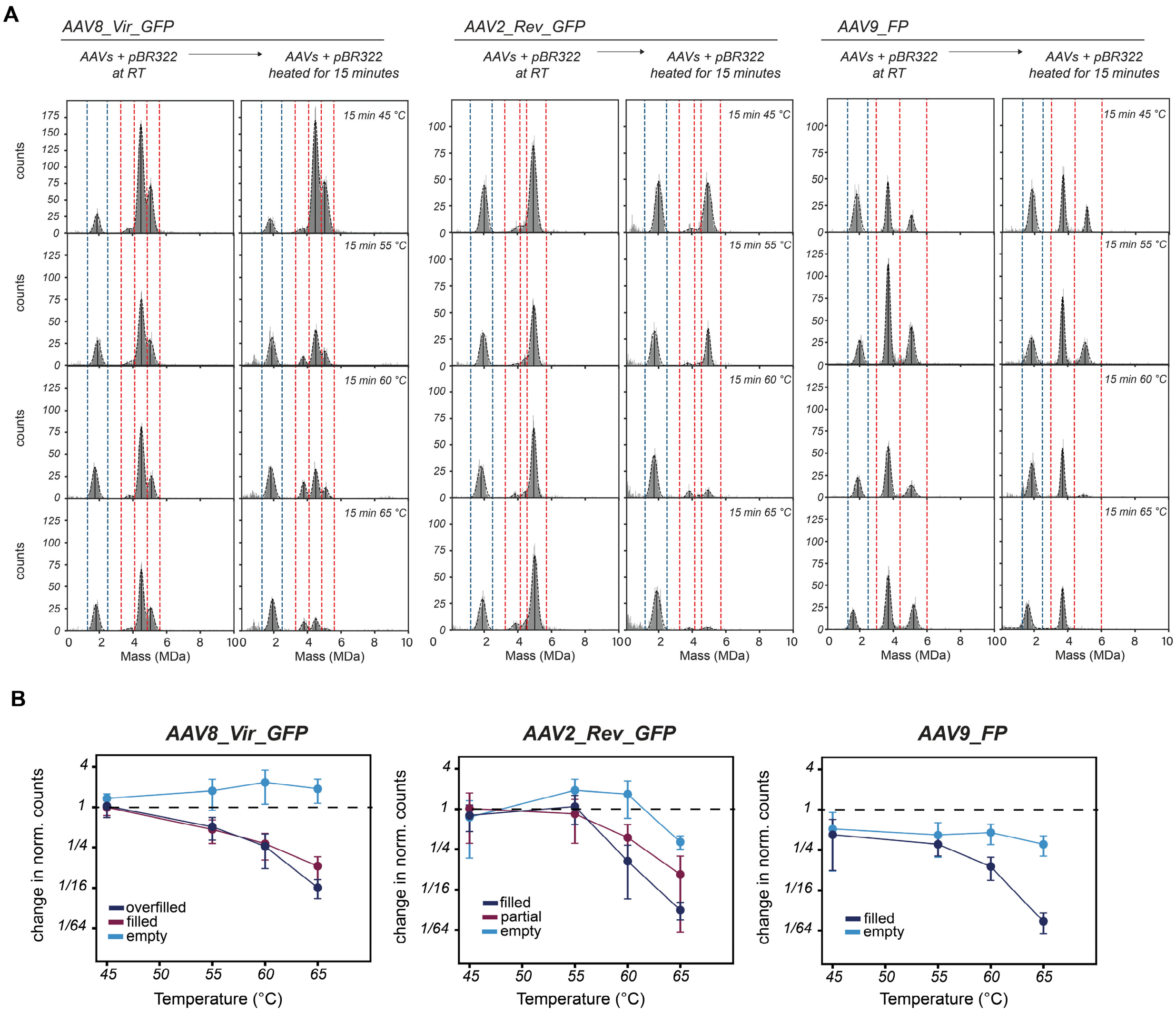


**Supplementary Figure 4:** ***Mass photometry quantification of thermal-stressed AAV8_Vir_GFP, AAV2_Rev_GFP and AAV9_FP together with the pBR322 plasmid standard.*** **A)** Illustrative mass histograms derived from MP measurements of the heat incubated AAV8_Vir_GFP, AAV2_Rev_GFP and AAV9_FP. AAVs were incubated at 45, 55, 60 and 65 °C together with the pBR322 reference plasmid for quantification and comparison to incubation at room temperature (RT). Gaussian fits of the AAVs and pBR322 are indicated by a black dashed line. The mass ranges that were used for quantification of the different AAV subpopulations are drawn by red dashed lines, the mass range used for pBR322 quantification is drawn by blue dashed lines (pBR322: 1.2 – 2.5 MDa, AAV8_Vir_GFP empty: 3.3 – 4.1 MDa, AAV8_Vir_GFP filled: 4.1 – 4.85 MDa, AAV8_Vir_GFP overfilled: 4.85 – 5.6 MDa, AAV2_Rev_GFP empty: 3.25 – 4.15 MDa, AAV2_Rev_GFP partially filled: 4.15 – 4.55 MDa, AAV2_Rev_GFP filled: 4.55 – 5.7 MDa, AAV9_FP empty: 3.0 – 4.4 MDa, AAV9_FP filled: 4.4 – 6.0 MDa). **B)** The heat induced loss of AAV8_Vir_GFP, AAV2_Rev_GFP and AAV9_FP was derived following normalization of the pBR322 signal before and after heating. Displayed are the relative changes in AAV subpopulations following incubation at room temperature compared to incubation at 45, 55, 60 and 65 °C. Error bars represent the standard deviation between the different repeats.

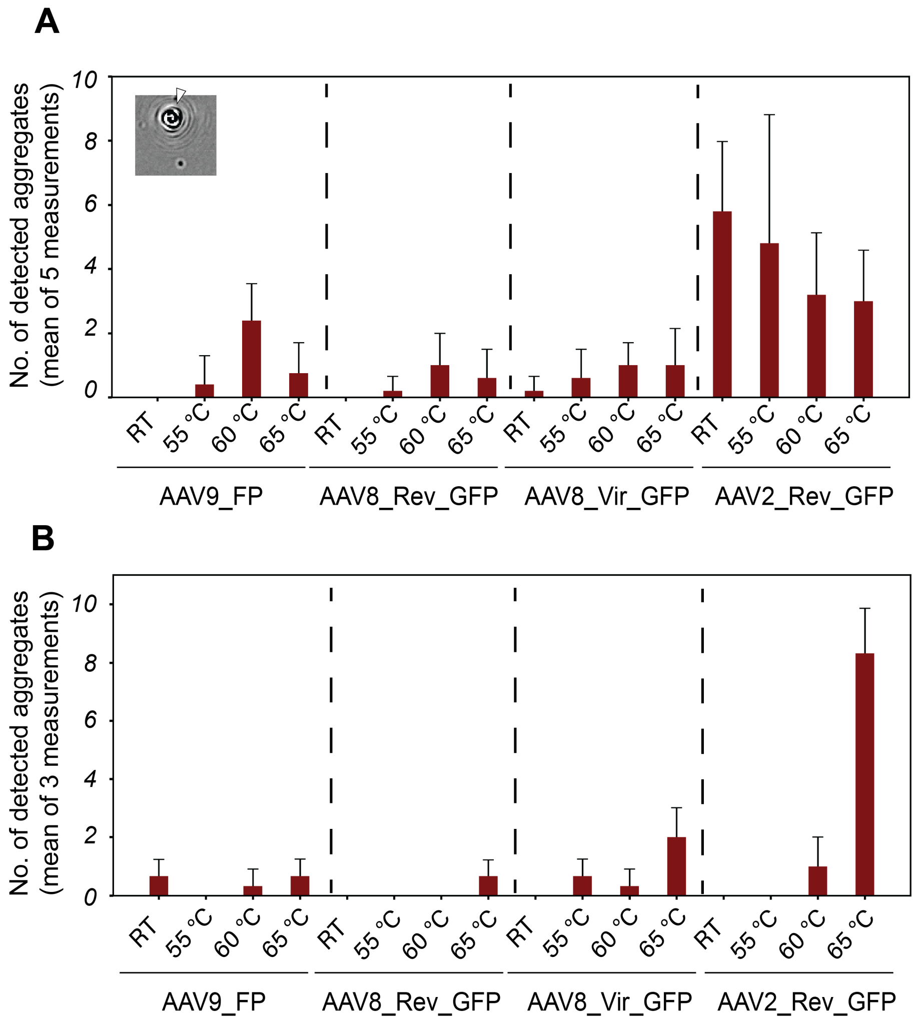


**Supplementary Figure 5:** ***Aggregation of AAVs following heating.*** Aggregates were identified by visual inspection of the MP recordings after incubation of 15 minutes at the indicated temperatures. **A)** Recordings of experiments from figure 3 and S4 that were performed on APTES slides were analyzed for aggregates. Within the AAV2_Rev_GFP sample considerably more aggregates could be seen compared to the other AAVs. Inset displays an example of an aggregate landing event. **B)** Recordings of experiments from figures 4 and 5 performed on glass slides were analyzed for aggregates. Here, AAV2_Rev_GFP showed prominent aggregation after heating at 65 °C. Error bars in both panels represent the standard deviation between the different repeats.

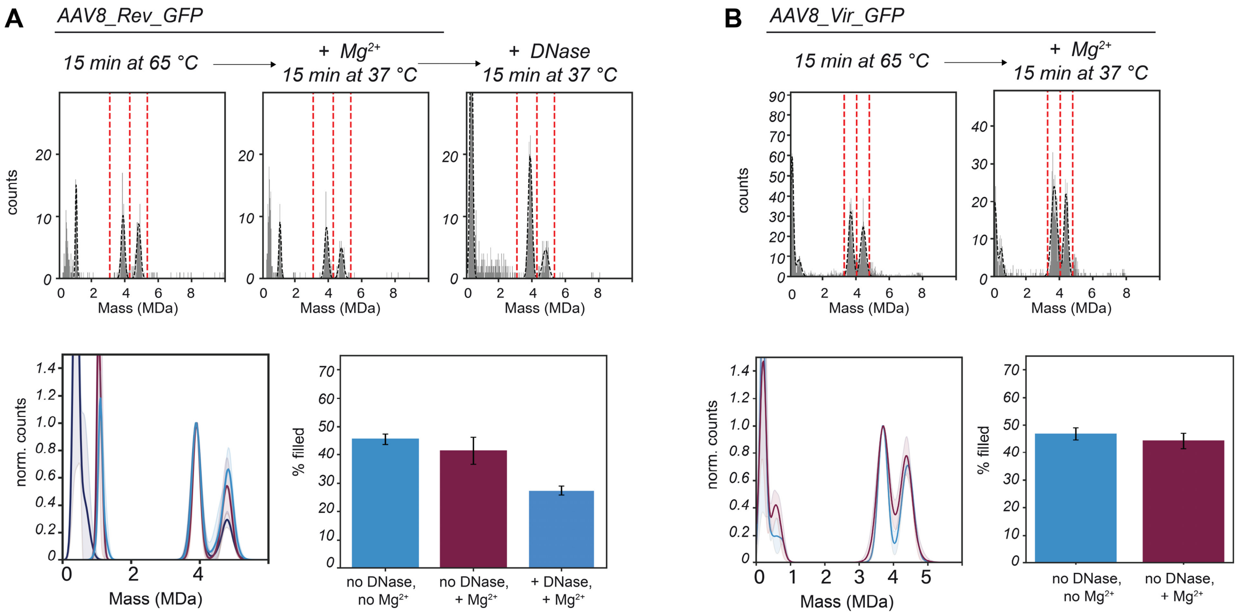


**Supplementary Figure 6:** ***no effect of $Mg^{2+}$ on the empty/filled distribution of AAV8 capsids.*** For both the AAV8 capsids (AAV8_Rev_GFP and AAV8_Vir_GFP) the change in empty and filled capsids were assessed by MP following addition of $Mg^{2+}$. No effect could be detected of $Mg^{2+}$ by itself. **A)** After heating of AAV8_Rev_GFP at 65 °C for 15 minutes the amount in empty and filled capsids were quantified within the specified range (indicated by red dashed lines). Three repeats were aligned and normalized on the most abundant AAV peak. The standard deviation is shown by semi-transparent bands. A bar-plot is shown with the percentage of filled AAVs. Only after addition of DNase a major drop in filled particles can be seen. **B)** The same approach was done for AAV8_Vir_GFP, where also no difference could be seen for addition of $Mg^{2+}$ alone.
